# Does ivermectin treatment for endemic hookworm infection alter the gut microbiota of endangered Australian sea lion pups?

**DOI:** 10.1101/2022.09.14.508058

**Authors:** Mariel Fulham, Michelle Power, Rachael Gray

## Abstract

The gut microbiota is essential for the development and maintenance of the hosts’ immune system, and disturbances can impact host health. This study aimed to determine if topical ivermectin treatment for endemic hookworm (*Uncinaria sanguinis*) infection in Australian sea lion (*Neophoca cinerea*) pups causes gut microbial changes. The gut microbiota was characterised for untreated (control) (n=23) and treated (n=23) pups sampled during the 2019 and 2020/21 breeding seasons at Seal Bay, Kangaroo Island. Samples were collected pre- and post-treatment on up to four occasions. The gut microbiota of both untreated (control) and treated pups was dominated by five bacterial phyla, Fusobacteria, Firmicutes, Proteobacteria, Actinobacteria and Bacteroides. There was a significant difference in alpha diversity between treatment groups in 2020/21 (p = 0.008), with greater diversity in treated pups. Modelling the impact of host factors on beta diversity revealed that pup ID accounted for most of the variation with pup ID, age and capture being the only significant contributors to microbial variation (p < 0.05). There were no statistically significant differences in microbial composition between treatment groups in both breeding seasons, indicating that ivermectin treatment did not alter microbial composition. To our knowledge, this is the first study to consider the impact of parasitic treatment on overall diversity and composition of the gut microbiota. Importantly, the lack of compositional changes in the gut microbiota with topical treatment support the utility of topical ivermectin as a safe and minimally invasive management strategy to enhance pup survival in this endangered species.

**Importance:** Disturbances to the gut microbiota in early life stages can have life-long impacts on host health. Australian sea lions are endangered and declining, and pups are endemically infected with hookworm (*Uncinaria sanguinis*) which contributes to pup mortality. Treatment with topical ivermectin has been shown to effectively eliminate hookworm infection and to improve pup health, but the impact on the gut microbiota was previously unknown, representing a key knowledge gap. The results from this study show that topical ivermectin treatment does not alter the gut microbiota of Australian sea lion pups, indicating that it is a safe and minimally invasive treatment that can aid in disease mitigation and conservation of this endangered species.

## 1. Introduction

The Australian sea lion is Australia’s only endemic pinniped species and has a highly fragmented population, breeding across 80 colonies which extend from the Houtman Abrolhos in Western Australia to The Pages Islands in South Australia (1, 2). The Australian sea lion population has undergone continual decline since commercial harvesting in the 18^th^ and 19^th^ centuries (3). It is currently listed as endangered on both the IUCN Red List (4) and the Environmental Protection and Biodiversity Conservation Act (EPBC) (5) with approximately 10,000 free-ranging individuals remaining (6), making it one of the rarest pinniped species in the world. The reasons for this decline are multifactorial, with fisheries interactions (7), entanglement in marine debris (8, 9), anthropogenic pollution (antibiotic resistant and human-associated bacteria, chemicals, and heavy metal contaminants) (10, 11, Taylor et al., in review) and disease (13, 14) identified as contributing factors.

At two of the largest Australian sea lion breeding colonies, Seal Bay Conservation Park on Kangaroo Island and Dangerous Reef in the Spencer Gulf, high levels of pup mortality have been reported, with rates of up to 41.8% and 44.6%, respectively (15, 16). Pup mortality within these colonies have been attributed to starvation, conspecific trauma, and stillbirths (17). Pups at both of these colonies are endemically infected with *Uncinaria sanguinis*, a haematophagous nematode (14) that causes localised intestinal inflammation, anaemia and hypoproteinaemia (13). *Uncinaria sanguinis* infection is typically patent for 2-3 months, and after clearance of infection, pups do not become re-infected (14). Given the continual population decline and high rates of pup mortality, recent conservation management for this species has included mitigation of pup mortality through hookworm treatment (18, 19). Lindsay et al. (2021) (19) determined that topical ivermectin treatment was 96.5% effective at eliminating hookworm with improved haematological parameters and increased bodyweight in treated compared to untreated pups. Hookworm infection is also prevalent in other pinniped species, contributing to differing levels of clinical disease and mortality in northern fur seal (*Callorhinus ursinus*) (20, 21), California sea lion (*Zalophalus californianus*) (22), South American fur seal (*Arctocephalus australis*) (23, 24) and New Zealand sea lion (*Phocarctos hookeri*) (25, 26) pups. Ivermectin has been administered to New Zealand sea lion, northern fur seal, and South American fur seal pups to reduce hookworm-associated mortality, and in all studies treatment resulted in hookworm clearance and improved pup growth and survival (21, 23, 26, 27). However, previous studies have not considered the potential consequences of treatment intervention on the composition of the gut microbiota in pinniped pups.

The gut microbiota of numerous pinniped species has been characterised (28–34), including in adult Australian sea lions (35). In most of these pinniped species, the gut microbiota is usually dominated by the *Firmicutes, Bacteroidetes*, *Actinobacteria* and *Proteobacteria* phyla, with the relative abundance differing between species and sites (29, 31, 34, 35). However, many of these studies have been limited to captive or adult pinnipeds, with the gut microbiota of pups only characterised in Juan Fernández fur seals (*Arctocephalus philippii*) (33), spotted seals (*Phoca larga*) (30), Australian fur seals (*Arctocephalus pusillus doriferus*) (32), and southern and northern elephant seals (*Mirounga leonina* and *Mirounga angustirostris*) (28, 31)

The mammalian gastrointestinal (GI) tract is home to over 100 trillion microorganisms, collectively termed the microbiota (36–38). Mammalian hosts and their microbiota have co-evolved over millions of years to establish a mutualistic relationship, and the importance of this relationship for host health has been established in recent decades (39, 40). Commensal bacteria contribute to digestion and nutrient provision (41), synthesise essential vitamins and minerals (42), protect against pathogen colonisation through colonisation resistance (CR) 1. (43) and are essential for the development and regulation of the immune system (40, 44).

The impact of intestinal parasites, such as helminths, on the composition of the gut microbiota remains relatively unknown, varying between host and parasite species (45–47). Helminth infections in laboratory mice can result in substantial shifts in the gut microbiota composition (48–51), which could be a consequence of multiple factors, including helminth secretory products (47). As intestinal helminths can both directly and indirectly interact with the gut microbiota, the removal of parasites via antiparasitic administration also has the potential to alter the composition of the microbial community (52, 53). In humans and mice, the removal of intestinal helminths has varying impacts on the gut microbiota; outcomes of parasite treatment range from no associated change (54), minimal to moderate change in microbiota composition (54, 55), to a shift in the composition of the microbiota to resemble individuals without helminth infection (56). In wildlife species, treatment with antiparasiticides has been associated with alterations to the faecal gut microbiota and metabolic profile (57, 58). Given the cross-talk between the gut microbiota and the immune system, it is likely that the modulation of the microbial community by intestinal helminths can have both direct and indirect impacts on the hosts’ immune system and immune responses (47, 53, 59). Given intestinal parasitism is prevalent in many free-ranging wildlife species (60), it is crucial to understand how parasitic infection (or removal thereof) influences gut microbial composition.

Just as microbes in the gut environment have evolved with their hosts, parasites can evolve with their hosts (61), and it is likely *U. sanguinis* has evolved with its host species. Furthermore, interactions between the host, intestinal helminths and the gut microbiota are likely to be complex and multidirectional (62). While ivermectin treatment has been shown to improve Australian sea lion pup health parameters in the short term (18, 19), the impact of removal of *U. sanguinis* on the gut microbial community, and potentially on pup health notwithstanding the improvements associated with mitigating hookworm disease, is unknown. The gut microbiota becomes established during early mammalian life stages, with disruptions or alterations potentially impacting the functional capacity of the microbiome, the development of the immune system (40, 63), altering the microbial composition and promoting colonisation of pathogenic bacteria, and most importantly, has the potential to influence lifelong host health and disease status (64, 65). Given the importance of the gut microbiota and its influence on development during early life stages, investigating whether treatment intervention and parasite elimination alters the gut microbiota of Australian sea lion pups is critical for understanding any potential consequences.

In order to establish the safety of a topical antiparasitic treatment as a potential management strategy to assist in the recovery of the Australian sea lion, the impact of treatment on the gut microbial community must be understood. To this end, this study aims to characterise and monitor the microbial composition of the gut microbiota of both treated (and subsequently hookworm-free) and untreated (control) Australian sea lion pups.

## 2. Methods

### 2.1. Study site and sample collection

Faecal samples were collected from neonatal Australian sea lion pups at Seal Bay Conservation Park on Kangaroo Island, South Australia. Pups were sampled during the 2019 winter (*n*=160) and 2020/21 summer (*n*=184) breeding seasons as part of an ivermectin treatment trial (Lindsay et al. unpublished).

In brief, pups were captured on up to four occasions, at approximately four-week intervals. Pups were captured by hand and physically restrained in a ventilated canvas bag designed specifically for pinniped pups. During initial capture (capture 1 - the time point prior to treatment administration), pups were assigned to a treated (and subsequently hookworm-free, herein referred to as treated) or untreated (control) group based on a randomised number chart, generated using Microsoft Excel. Morphometric data was collected from each pup during each capture event including bodyweight (kg), standard length (cm, measured from tip of the nose to tip of the tail), sex and body condition (poor, fair-thin, good, excellent). The initial capture also included a unique ‘hair cut’ on the dorsolumbar pelage and application of commercial hair dye (Schwarzkopf Nordic Blonde, Henkel Australia, Melbourne, Australia) to facilitate individual pup identification for recapture. Faecal swabs were collected via insertion of a sterile swab (Copan, Brescia, Italy) within a lubricated sheath into the rectum. Swabs were then subsampled into Sterile FecalSwab™ tubes (Copan, Brescia, Italy) and stored at 4°C for up to 2 months, followed by storage at -20°C for up to 2 months, and then stored at -80°C. Blood samples were collected from the brachial vein as previously described (10, 66). Up to 1mL of blood was transferred into a tube containing ethylenediaminetetraacetic acid (EDTA) (Sarstedt, Nümbrecht, Germany) for haematological analysis. Blood samples were stored at 4°C and processed within 10 hours of collection. Sampling of Australian sea lion pups was approved by the Animal Ethics Committee at the University of Sydney (Protocol Number 2017/1260).

Of the total number of pups sampled in 2019 and 2020/21, a subset of pups (*n*=46) that had been captured on four occasions were randomly selected from both the untreated (control) and treated groups by assigning a random number (Microsoft Excel for Mac v16.61.1) to each pup ID for inclusion in the microbial study.

### 2.2. Hookworm infection status

After collection, faecal swabs were kept at 4°C prior to storage at -20°C. Samples were processed within 14 days of collection. The hookworm status of each pup was determined following methods described by Lindsay et al. (2021) (19). In summary, faecal material was transferred onto a clean glass slide and examined via light microscopy for the presence of *U. sanguinis* eggs. In samples where no eggs were detected, two additional smears were evaluated to confirm negative status.

### 2.3. Haematological analysis

Blood samples were processed following methodology described by Lindsay et al. (2021) (19). In brief, the packed cell volume (PCV; L/L) was measured using the microhaematocrit method and total plasma protein (TPP; g/L) was estimated using a hand-held refractometer (Reichert TS Meter, Cambridge Instruments, Buffalo, USA). Blood smears were prepared in duplicate and fixed in 100% methanol (Chem-Supply Pty Ltd, Port Adelaide, South Australia) for 4 minutes. An aliquot (200µL) of anticoagulated EDTA blood was transferred to a separate tube for preservation with an equal aliquot (200µL) of Streck Cell preservative (Streck, Omaha, USA) and stored at 4°C prior to further analysis. Preserved samples were analysed on an automated haematology analyser (Sysmex XT-2000iV, Sysmex, Kobe, Japan) at the Veterinary Pathology Diagnostic Service, Sydney School of Veterinary Science, The University of Sydney, within 2-8 days of sample collection. From automated haematology analysis, the total erythrocyte count (x10^12^/L), haemoglobin concentration (g/L), mean cell volume (fL), mean cell haemoglobin concentration (g/L), platelet count (x10^9^/L) and total nucleated (leukocyte) cell count (TNCC, x10^9^/L) were determined. The differential leukocyte count was obtained by differentiating 100 leukocytes for every 10×10^9^/L TNCC to determine the absolute neutrophil, lymphocyte, eosinophil and monocyte counts (x10^9^/L) by multiplying the percentage of each leukocyte by the TNCC. Nucleated red cell (nRBC) count was determined by counting the number of nucleated red blood cells per 100 leukocytes.

### 2.4. Age determination

As part of an ongoing monitoring program at Seal Bay Conservation Park, the breeding colony is frequently monitored during the breeding season, allowing birth dates to be recorded (67). Date of birth for pups was provided by the Department of Environment and Water (DEW), South Australia and the South Australian Research and Development Institute (SARDI), as part of ongoing monitoring within each breeding season. For the 2020/21 breeding season, all pups included in the study had known dates of birth; age at each capture was calculated based on known birth date. For the 2019 breeding season, birth dates were only known for a subset of pups (*n*=13). For the remaining pups (*n*=10) sampled in 2019, a regression analysis was utilised to estimate pup age, following the approach of Bradshaw et al. (2000) (68). The fitted age model for the 2019 data is:

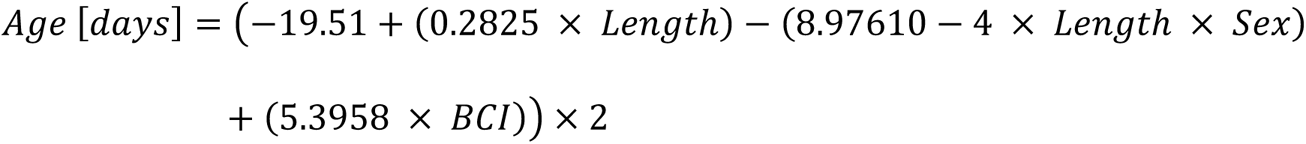

where Sex has the value of 1 for males and 0 for females, and BCI (body condition index) is calculated based on a regression of length on weight (69).

### 2.5. DNA extraction and PCR amplification

From the randomised subset of pups, DNA was extracted from a total of 184 faecal swabs (*n*=92 from 2019 and *n*=92 from 2020/21). DNA was also extracted from two faecal samples collected from *N. cinerea* pups in 2019 to be used as control samples to account for variation between sequencing runs.

Faecal DNA was extracted from FecalSwab^TM^ media (200µL) using the ISOLATE II Fecal DNA Kit (Bioline, Sydney, Australia) following the manufacturer’s protocol. Faecal DNA was tested for PCR competency by a 16S PCR using methods described by Fulham et al. (2020) (10) using forward primer 27F and reverse primer 1492R (70). Amplicons were resolved using gel electrophoresis (2% agarose w/v) with SYBR safe gel stain (Invitrogen, Sydney, Australia) and conducted at 100V for 30 min with product size approximated using HyperLadder II 50bp DNA marker (Bioline, Sydney, Australia).

### 2.6. 16S rRNA sequencing

Prior to sequencing, the concentration of DNA (ng/µL) and purity of nucleic acids in all faecal DNA was tested using a NanoDrop® ND-1000 UV-Vis Spectrophotometer (Thermo Fisher Scientific, Massachusetts, United States). DNA was submitted to the Ramaciotti Centre for Genomics (University of New South Wales, Sydney, Australia) for 16SV1-3 amplicon sequencing with an Illumina Miseq v3 2×300 bp sequencing kit using primers 27F and 519R, producing a ∼530bp fragment Lane (1991) (70).

### 2.7. Analysis of sequences and taxonomic classification

Demultiplexed paired-end sequences were analysed using Quantitative Insights Into Microbial Ecology 2 (QIIME 2) version 2022.2 software (71). The DADA2 (Divisive Amplicon Denoising) plugin (72) in QIIME 2 was utilised to trim and filter sequences for quality including the removal of primers, denoising and chimera removal. Sequences were clustered into amplicon sequence variants (ASVs), also known as exact sequence variants, following denoising. Operational taxonomic units (OTUs) are also frequently used to define the gut microbiota and clusters sequences either based on the observed sequences or using a reference database (73). Recently, more focus has been on ASVs as they are more sensitive and specific than OTUs, can be reused across studies, and are reproducible for future data sets (74).

Taxonomies were assigned to each ASV using a 16S rRNA V1-V3 classifier trained against the frequently updated SILVA database (release 138) (75–77) using the q2-feature-classifier plugin in QIIME 2 (78). Low abundance ASVs that were not present in at least two samples with a total read count below 20 were filtered out and removed prior to further analysis.

### 2.7. Statistical analyses

All statistical analyses were conducted in either QIIME 2 or RStudio (v2022.2.3+492, Boston Massachusetts). Data was exported from QIIME 2 into R using the QIIME2R package (79). Using both QIIME 2 and the vegan package (80), alpha diversity rarefaction curves were generated to filter samples based on sufficient sampling depth and data was rarefied with the threshold of 5000 reads used as the cut-off for samples collected in both 2019 and 2020/21. Significance was determined when p > 0.05 for all statistical tests.

For analysis of pup morphometric and health data, pups were grouped into five weight groups (kg; 6.0-9.9, 10.0-13.9, 14.0-17.9, 18.0-21.9, 22.0-25.9), six standard length groups (cm; 65.0-69.9, 70.0-74.9, 75.0-79.9, 80.0-84.9, 85.0-89.0, 90.0-94.9) and seven age groups (months; 0.0-0.5, 0.5-1.0, 1.0-2.0, 2.0-3.0, 3.0-4.0, 4.0-5.0, 5.0-6.0).

#### 2.7.1 Alpha diversity analysis

The alpha (within-sample) diversity was estimated using the phyloseq package (81). Richness, the total number of bacterial species present, and diversity, the amount of individual bacteria from each species identified, were measured using two metrics: Chao1, which estimates the richness in a sample through the estimation of the total number of species present (82); and the Shannon-Wiener index, which estimates diversity based on richness and abundance (83). A Shapiro-Wilk test was used to test for data normality. Wilcoxon’s Rank Sum Tests were employed to estimate the statistical differences in alpha diversity metrics due to treatment group (untreated [control] and treated) and breeding seasons. Statistical differences in alpha diversity pre-treatment (capture 1) and post-treatment (capture 2) for both treatment groups were also tested using Wilcoxon’s Rank Sum Test.

To examine the correlations between host factors and alpha diversity, four linear mixed models (LMMs) were fitted using the lme4 package v1.1-25 (84). The first two models were conducted when analysing entire datasets within a season. The first model included treatment group, capture (1–4), pup sex, weight (kg), standard length (cm) and age (months) as predictors and pup ID as a random effect, and the alpha diversity metric (Chao1 or Shannon-Wiener Index) as the response variable. In the second model, treatment group, hookworm status, age (months), TPP, TNCC, leukocyte counts (total neutrophil, total monocyte, total lymphocyte and total eosinophil; x10^9^/L) were predictors, pup ID was a random effect, and the alpha diversity metric was the response variable. Models three and four tested the influence of host factors within treatment groups and were the same as models one and two, respectively, with treatment group excluded.

#### 2.7.2. Beta diversity analysis

The variation between samples (beta diversity) was analysed using Bray-Curtis dissimilarity calculations using rarefied data based on the abundance of ASVs and using principal coordinate analysis (PCoA) plots. Using the adonis function in the vegan package (80), a permutational analysis of variance (PERMANOVA) was employed to test the statistical differences in ASV abundances due to treatment group, hookworm status, weight and standard length groups, sex, capture, and pup age, and computed with 999 permutations. Four PERMANOVA models were fitted; the first included pup ID, hookworm status and treatment group; the second model included pup ID, treatment group, capture, and age; the third model included treatment group, pup ID, weight and standard length groups and pup sex; and the final model included pup ID, treatment group, TPP, TNCC and leukocyte counts (neutrophils, lymphocytes, monocytes and eosinophils). The second model was compared *post hoc* to account for effects of repeated measures. The assumption of multivariate homogeneity was met across all models for PERMANOVA tests (p > 0.05).

Wilcoxon’s Rank Sum Test was used to test for significant differences in the relative abundance of bacterial phyla and families between treatment groups and within and between breeding seasons. Finally, an analysis of composition of microbiomes (ANCOM) was utilised to test for differences in differentially abundant ASVs between treatment groups (85), identifying any bacterial families that had the highest contribution to any dissimilarities observed in the gut microbial composition of untreated (control) and treated groups.

## 3. Results

Randomisation of the sampled pups resulted in the sequencing of samples from *n*=12 untreated (control) and *n*=11 treated pups from the 2019 breeding season, and *n*=11 untreated (control) and *n*=12 treated pups from 2020/21 breeding season. Treatment with ivermectin resulted in 100% elimination of hookworm infection in the *n*=46 Australian sea lion pups included in this study (Lindsay et al. unpublished). The hypervariable V1 to V3 region of the 16s rRNA gene was sequenced in a total of 184 samples collected in 2019 and 2020/21, resulting in 12,078,034 and 8,439,797 sequences, respectively. After filtering steps and removal of low quality (*n*=32) and control samples (*n*=2), 75 samples from 2019 and 77 samples from 2020/21 were imported into RStudio for analysis. The filtering of low abundance ASVs resulted in the clustering of 621 and 512 ASVs in the 2019 and 2020/21 datasets, respectively.

### 3.1.1 Gut microbiota composition of untreated (control) and treated Australian sea lion pups during the 2019 breeding season

Five bacterial phyla, *Fusobacteria*, *Firmicutes, Actinobacteria*, *Proteobacteria* and *Bacteroidetes*, were present in untreated (control) and treated Australian sea lion pups (Figure 1). The majority of ASVs in both untreated (control) and treated pups were assigned to the *Fusobacteria* phylum (Table 1). In both groups, *Fusobacteria* had the highest average relative abundance (average relative abundance ± standard deviation = 55.4% ± 27.5 in untreated (control) and 61.3% ± 27.1 in treated), followed by *Firmicutes* (27.1% ± 12.9 and 20.5% ± 9.2), *Proteobacteria* (10.3% ± 4.3 and 9.2% ± 5.7), *Actinobacteria* (6.3% ± 3.3 and 6.5% ± 3.7) and *Bacteroidetes* (0.1% ± 0.1 in both treatment groups). There was no significant difference in the average relative abundance of each bacterial phyla pre- or post-treatment (capture 1-capture 2; p > 0.05) in both treatment groups, or across captures 1 – 4 (p > 0.05). In both untreated (control) and treated pups, the relative abundance of *Fusobacteria* remained stable across all captures, while the abundance of *Firmicutes* decreased from capture 1 to capture 4 (Figure 1). The abundance of *Actinobacteria* increased after the second capture and then decreased in captures 3 and 4 in pups from both treatment groups, with the exception of one pup (pup-19-141) in the untreated (control) group with a much greater abundance of *Actinobacteria* after capture 4 (Figure 1).

**Figure 1.**
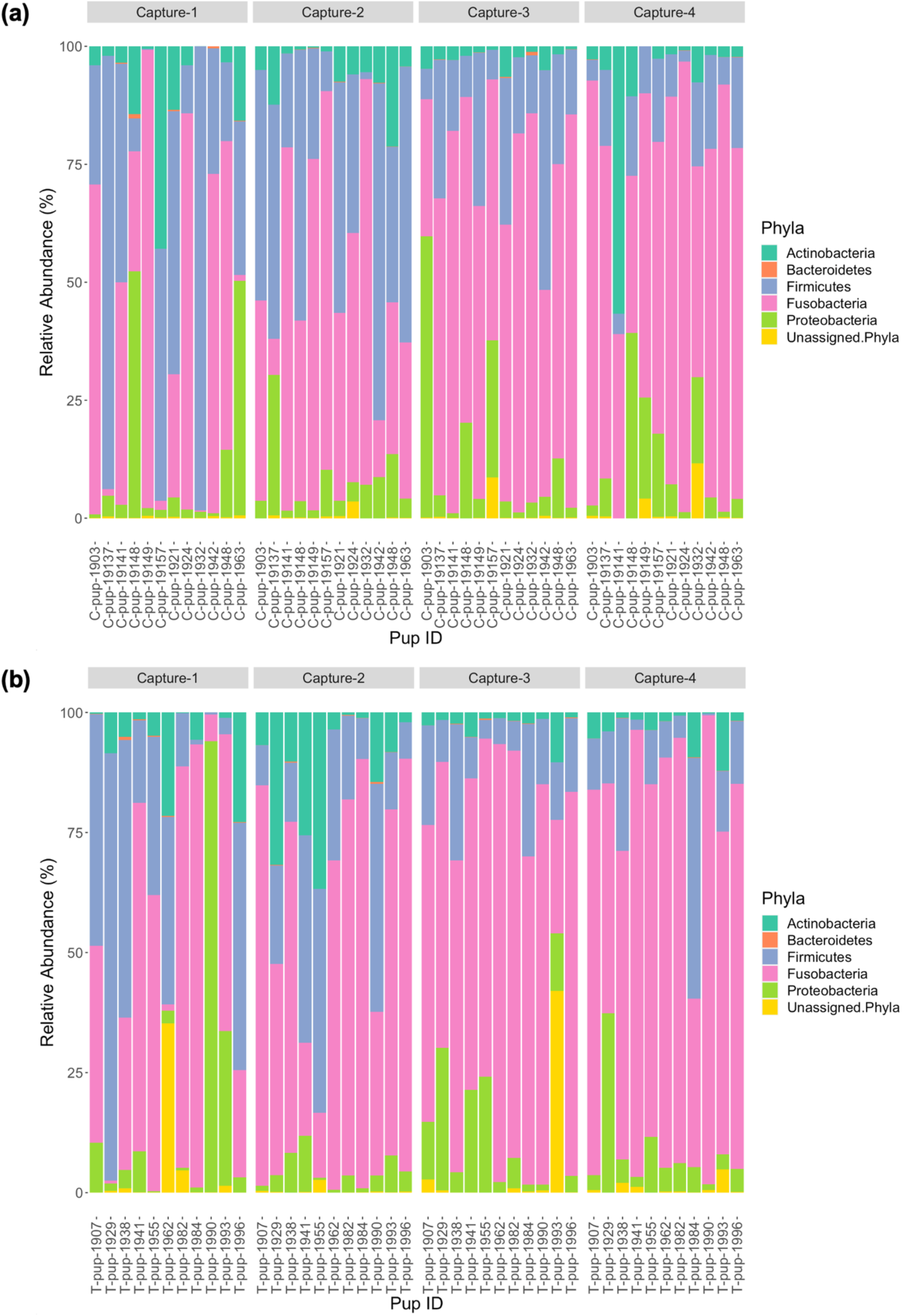
Relative abundance (%) of bacterial phyla in (a) untreated [control] and (b) treated Australian sea lion pups across four captures during the 2019 breeding season.

**Table 1.** Abundance of ASVs in untreated (control) and treated pups in captures 2-4, present in at least 90% of samples in each treatment group. The abundance of ASVs from first captures were excluded from both treatment groups to reflect any changes in abundance after treatment was administered.

| ASV | Phylum | Class | Family | Genus | Treatment Group | Abundance (%) | Year |
| --- | --- | --- | --- | --- | --- | --- | --- |
| 9ea04ed3882733d04922cb403e743965 | Fusobacteria | Fusobacteriia | Fusobacteriaceae | Fusobacterium | Untreated (control) | 6.1 | 2019 |
| 89f44d1e04224365282d4702046f1ce9 | Fusobacteria | Fusobacteriia | Fusobacteriaceae | Fusobacterium | Untreated (control) | 6.0 | 2019 |
| 147eb0d49c44569aa2f12c06fce289a0 | Fusobacteria | Fusobacteriia | Fusobacteriaceae | Fusobacterium | Untreated (control) | 4.7 | 2019 |
| 14ac82f71eee6441a2decab6f008b8e0 | Actinobacteria | Coriobacteriia | Coriobacteriaceae | Collinsella | Untreated (control) | 0.9 | 2019 |
| 147eb0d49c44569aa2f12c06fce289a0 | Fusobacteria | Fusobacteriia | Fusobacteriaceae | Fusobacterium | Treated | 8.8 | 2019 |
| 89f44d1e04224365282d4702046f1ce9 | Fusobacteria | Fusobacteriia | Fusobacteriaceae | Fusobacterium | Treated | 5.9 | 2019 |
| 9ea04ed3882733d04922cb403e743965 | Fusobacteria | Fusobacteriia | Fusobacteriaceae | Fusobacterium | Treated | 4.9 | 2019 |
| 89f44d1e04224365282d4702046f1ce9 | Fusobacteria | Fusobacteriia | Fusobacteriaceae | Fusobacterium | Untreated (control) | 3.5 | 2020/21 |
| 9ea04ed3882733d04922cb403e743965 | Fusobacteria | Fusobacteriia | Fusobacteriaceae | Fusobacterium | Untreated (control) | 3.1 | 2020/21 |
| 89f44d1e04224365282d4702046f1ce9 | Fusobacteria | Fusobacteriia | Fusobacteriaceae | Fusobacterium | Treated | 6.1 | 2020/21 |
| 9ea04ed3882733d04922cb403e743965 | Fusobacteria | Fusobacteriia | Fusobacteriaceae | Fusobacterium | Treated | 3.8 | 2020/21 |
| d908d12d31a00d235b622945720d56bb | Firmicutes | Clostridia | Lachnospiraceae | Blautia | Treated | 1.0 | 2020/21 |

In both untreated (control) and treated pups, *Fusobacteriaceae* was the most dominant bacterial family. In untreated (control) pups, ASVs were assigned to five families from four bacterial phyla. The *Fusobacteriaceae* family occurred at the highest average relative abundance (54.1% ± 27.7), followed by families from the *Firmicutes* phyla, *Clostridiaceae* (17.8% ± 7.1) and *Ruminococcaceae* (7.4% ± 6.8), *Coriobacteriaceae* (4.4% ± 3.0) from the *Actinobacteria* phyla and *Succinivibrionaceae* (2.7% ± 1.7) from the *Proteobacteria* phyla. Seven bacterial families from four bacterial phyla were identified in treated pups, with *Fusobacteriaceae* being similarly dominant (55.2% ± 20.7). The *Clostridiaceae* family was also the second most abundant in treated pups (13.5% ± 9.4), followed by *Coriobacteriaceae* (5.7% ± 3.5), *Ruminococcaceae* (5.7% ± 4.1), *Succinivibrionaceae* (4.5% ± 2.3), *Rhodobacteraceae* (2.9% ± 1.1), a member of the *Proteobacteria* phyla, and *Leptotrichiaceae* (2.2% ± 0.5) a family belonging to *Fusobacteria* (Figure 2). There was no significant difference in the average relative abundance of each family between treatment groups (p > 0.05). The *Rhodobacteraceae*, *Leptotrichiaceae* and *Succinivibrionaceae* families were only identified at an abundance ≥1% in ASVs in treated pups, with *Leptotrichiaceae* only present in treated pups after capture 4 (Figure 2).

**Figure 2.**
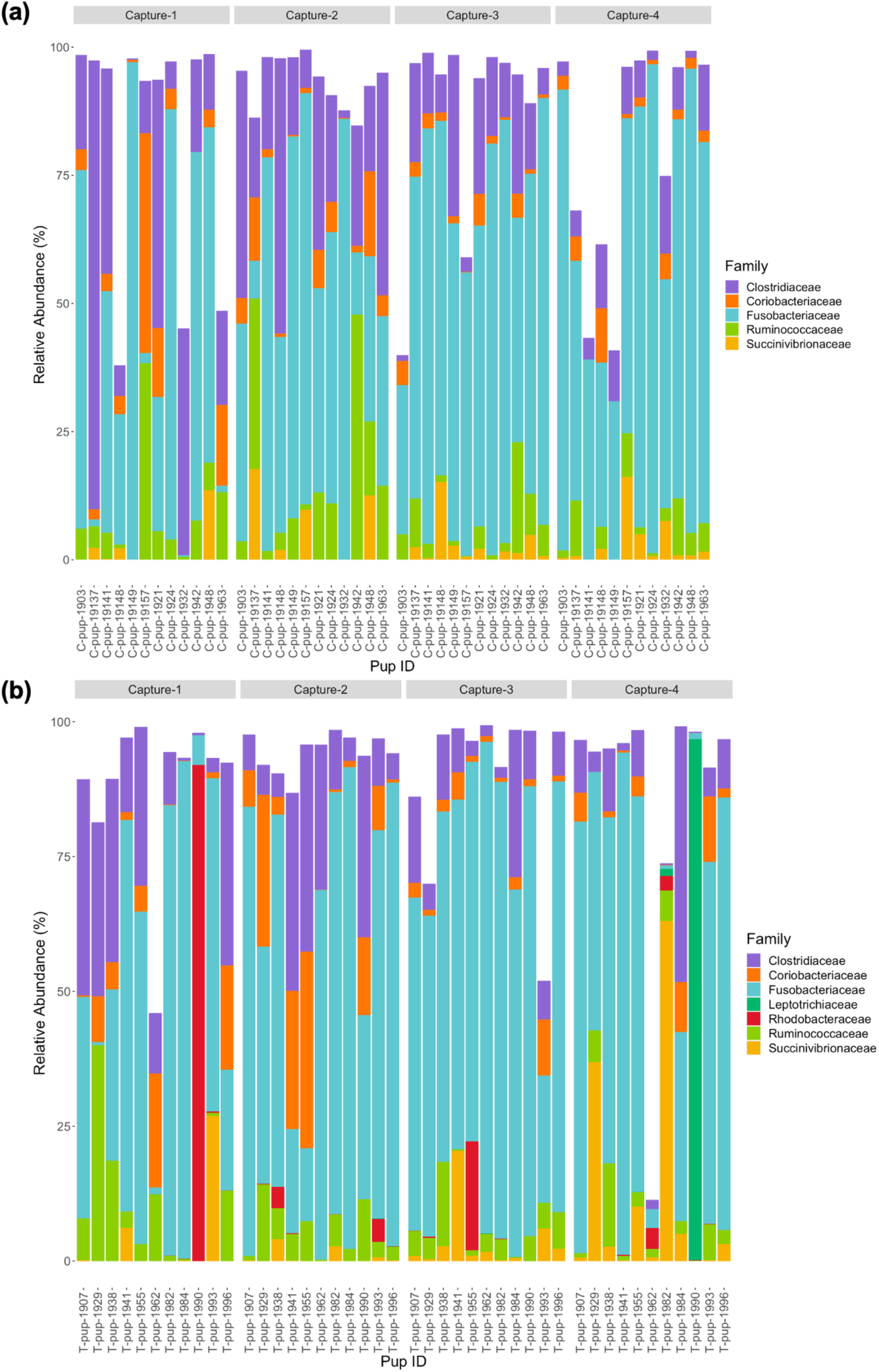
Relative abundance (%) of bacterial families in (a) untreated [control] and (b) treated Australian sea lion pups across all four captures during the 2019 breeding season. Only families that occurred at an abundance ≥ 1% in each treatment group are shown.

### 3.1.2. Gut microbiota composition of untreated (control) and treated Australian sea lion pups during the 2020/21 breeding season

In Australian sea lion pups sampled during the 2020/21 breeding season, five bacterial phyla were identified. The gut microbiota of both untreated (control) and treated pups was dominated by *Fusobacteria*, which had the highest average relative abundance (38.6% ± 23.2 in untreated [control] and 42.7% ± 20.9 in treated pups), followed by *Firmicutes* (36.1% ± 20.4 and 36.3% ± 20.3), *Proteobacteria* (16.7% ± 8.3 and 14.7% ± 6.3), *Bacteroidetes* (5.0% ± 1.7 and 4.5% ± 1.2) and *Actinobacteria* (3.4% ± 1.2) and 1.1% ± 0.3). The majority of ASVs in both treatment groups were assigned to the phylum *Fusobacteria* (Table 1). A small percentage of ASVs could not be assigned to a bacterial phylum in both groups, 0.2% in untreated (control) and 0.7% in treated pups. The average relative abundance of each bacterial phyla did not change when compared pre- and post-treatment in the treated group (p < 0.05). There was no significant difference in the average relative abundance of each bacterial phyla between treatment groups or across captures 1 - 4 (p > 0.05). The relative abundance of *Fusobacteria* and *Firmicutes* decreased in both treatment groups across the four captures while *Proteobacteria* and *Actinobacteria* increased (Figure 3). The abundance of *Bacteroidetes* was greatest in samples from capture 4 in both treatment groups (Figure 3).

**Figure 3.**
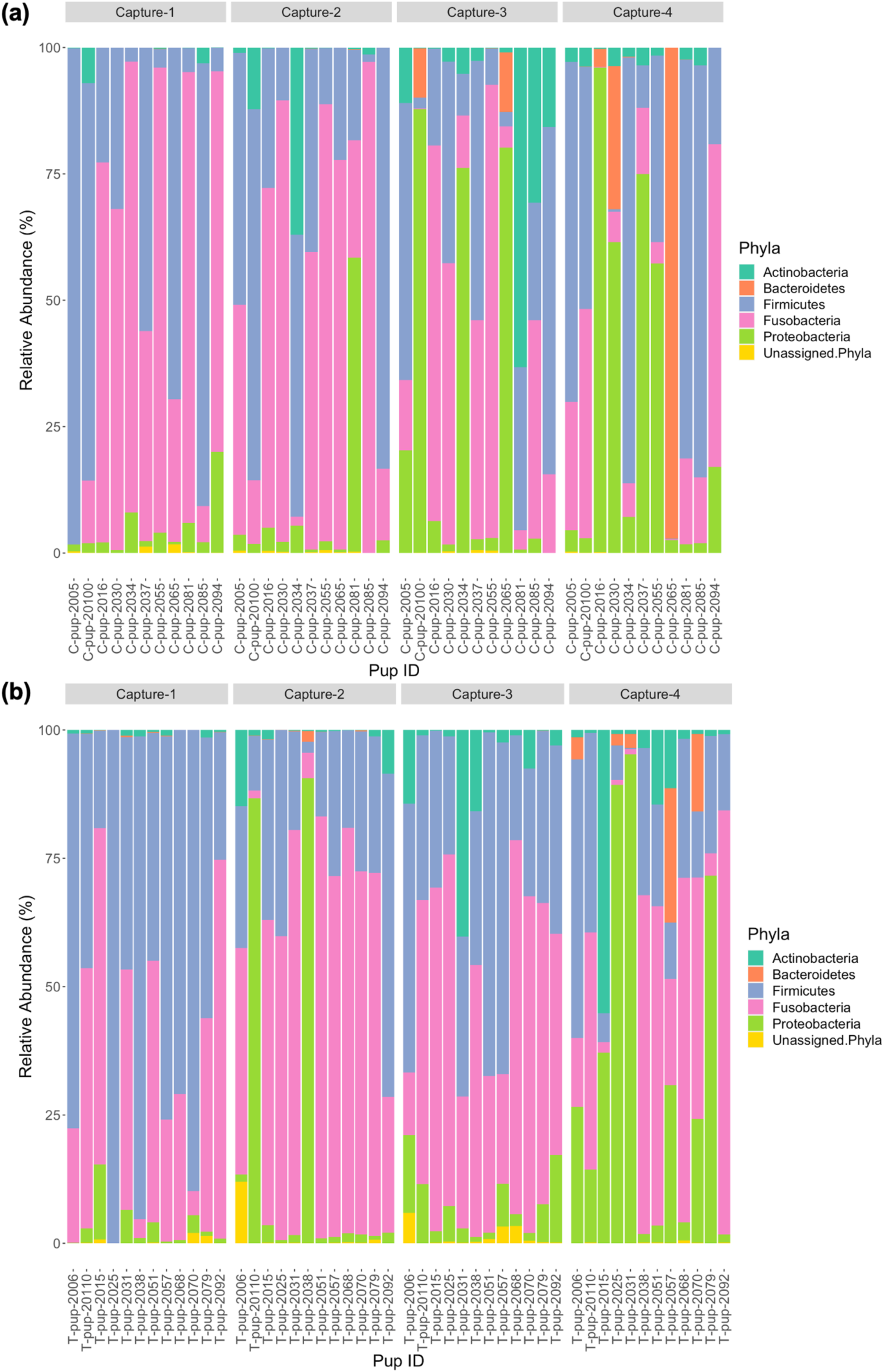
Relative abundance (%) of bacterial phyla in (a) untreated [control] and (b) treated Australian sea lion pups across the four captures during the 2020/21 breeding season.

At the family level, seven bacterial families were present in untreated (control) pups and five families were identified in treated pups that had a total abundance ≥ 1%. In untreated (control) pups, the average relative abundance was highest for the *Fusobacteriaceae* family (39.4% ± 23.9), followed by families belonging to the *Firmicutes* phyla, *Lachnospiraceae* (19.3% ± 7.6), *Clostridiaceae* (8.9% ± 3.1) and *Ruminococcacae* (5.0% ± 2.2); *Halomonadaceae* (7.5% ± 1.5) and *Rhizobioaceae* (3.1% ± 0.9) from the *Proteobacteria* phyla; and *Flavobacteriaceae* (3.2% ± 1.0) from the *Bacteroidetes* phyla. The five families assigned to reads from treated pups were similarly abundant, with *Fusobacteriaceae* being the most dominant (42.4% ± 27.1), followed by *Lachnospiraceae* (18.9% ± 10.2), *Clostridiaceae* (10.3% ± 7.7), *Halomonadaceae* (6.0% ± 5.8) and *Ruminococcaceae* (6.0% ± 5.0). While the abundance of *Fusobacteriaceae* decreased from capture 1 to capture 4 in both treatment groups, a more noticeable decrease in *Clostridiaceae* was apparent in treated pups after capture 1 (Figure 4). The *Flavobacteriaceae* and *Rhizobiaceae* families were only present in untreated (control) pups in captures 3 and 4 (Figure 4).

**Figure 4.**
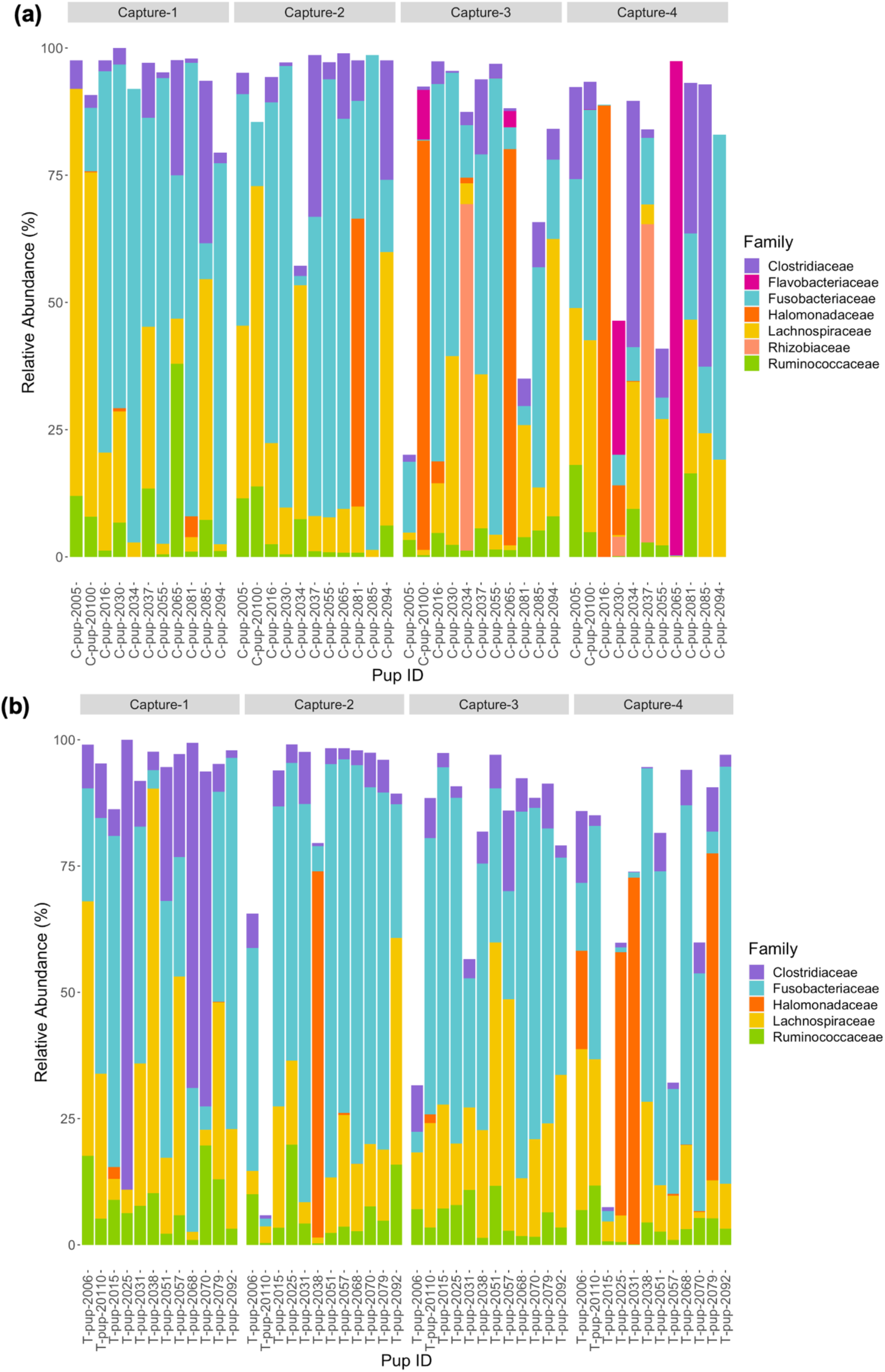
Relative abundance (%) of bacterial families in (a) untreated [control] and (b) treated Australian sea lion pups over four captures, sampled during the 2020/21 breeding season. Only families that occurred at an abundance ≥ 1% in each treatment group are shown.

### 3.1.3. Comparison of gut microbiota composition between breeding seasons

Five bacterial phyla were identified in both treated and untreated (control) pups across both breeding seasons. In both 2019 and 2020/21 respectively, the majority of ASV counts belonged to three phyla: *Fusobacteria* (53.5% and 39%), *Firmicutes* (22.1% and 33.8%) and *Proteobacteria* (9.6% and 14.4%). The *Actinobacteria* phyla was more dominant in 2019 (5.8% and 2.1%) while *Bacteroidetes* was more dominant in 2020/21 (0.1% and 4.4%). A small percent of ASVs, 1.6% and 0.43%, could not be assigned to a bacterial phylum in 2019 and 2020/21, respectively. There was a significant difference in the relative abundance of *Fusobacteria* (p < 0.001), *Firmicutes* (p = 0.0021), *Actinobacteria* (p < 0.0001) and ASVs that could not be assigned to a phylum (p = 0.001). The average relative abundance of *Fusobacteria* and *Actinobacteria* were higher in 2019 (58.6 ± 27.18 and 6.30 ± 3.43) compared to 2020/21 (41.4 ± 20.1 and 4.4 ± 2.3), while the opposite was observed for *Firmicutes* (23.8 ± 11.1 in 2019 and 33.8 ± 16.0 in 2020/21). There was no significant difference in the relative abundance of *Bacteroidetes* (p = 0.11) or *Proteobacteria* (p = 0.29) across the 2019 and 2020/21 breeding seasons.

When comparing the relative abundance of each phylum in untreated (control) pups across seasons, there was only a significant difference in *Fusobacteria* (p = 0.021) and *Actinobacteria* (p = 0.0023), with a greater relative abundance of both phyla in 2019. In treated pups, there was a significant difference between breeding seasons in the relative abundance of *Fusobacteria* (p < 0.01), *Firmicutes* (p < 0.01) and *Actinobacteria* (p < 0.01). The relative abundance of *Fusobacteria* and *Actinobacteria* was higher in treated pups in 2019, while *Firmicutes* occurred at a higher relative abundance in 2020/21.

Three families, *Fusobacteriaceae*, *Clostridiaceae* and *Ruminococcaceae* were present in pups in both breeding seasons. There was a significant difference in the relative abundance of *Fusobacteriaceae* (p = 0.001) and *Clostridiaceae* (p < 0.001) between the 2019 and 2020/21 breeding seasons. Similar significant differences were observed in untreated (control) pups between seasons in the *Fusobacteriaceae* (p = 0.026) and *Clostridiaceae* (p < 0.001) families, with both occurring at higher relative abundances in untreated (control) pups in 2019, compared to 2020/21. In treated pups, there was also a significant difference between breeding seasons in the relative abundance of *Fusobacteriaceae* (p = 0.018) and *Clostridiaceae* (p = 0.041), with the relative abundance of both families also being higher in Australian sea lion pups sampled during the 2019 breeding season compared to the 2020/21 season. For both treatment groups, there was no significant difference in the relative abundance of *Ruminococcaceae* (p > 0.05).

## 3.2. Alpha diversity

### 3.2.1. Comparison of alpha diversity between treatment groups and breeding seasons

Two measures of alpha diversity (Chao1 and Shannon-Wiener Index) were used to analyse differences in within-sample diversity between treatment groups and breeding seasons. To determine any immediate differences post-treatment, alpha diversity was compared between capture 1 and capture 2 for each cohort (2019 and 2020/21) as well as within each treatment group.

In the 2019 cohort, the average diversity, as indicated by the Shannon-Wiener index, was higher in the untreated (control) group compared to the treated group, while richness, measured through the Chao1 index, was higher in treated pups (Table 2, Figure 5). There was no significant difference in alpha diversity between treatment groups based on the Chao1 index (p = 0.157) and the Shannon-Wiener index (p = 0.159). There was also no significant difference in alpha diversity between capture 1 and capture 2 in the untreated (control) and treated groups based on both the Chao1 index (p = 0.318 and p = 0.558) and the Shannon-Wiener index (p = 1.00 and p = 0.644).

**Table 2.**
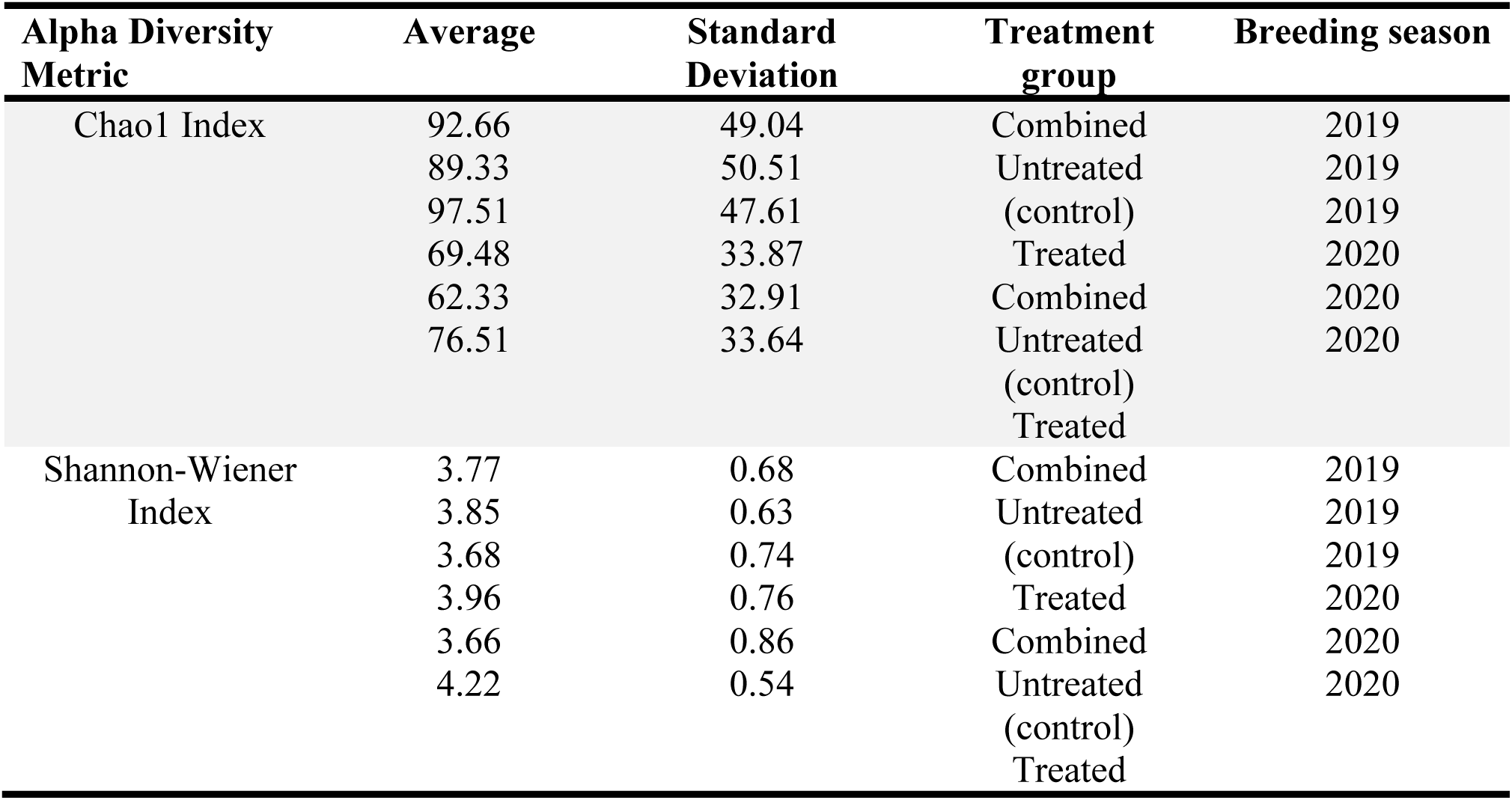
Mean Chao1 index and Shannon-Wiener indices for both untreated (control) and treated groups in both the 2019 and 2020/21 breeding seasons.

**Figure 5.**
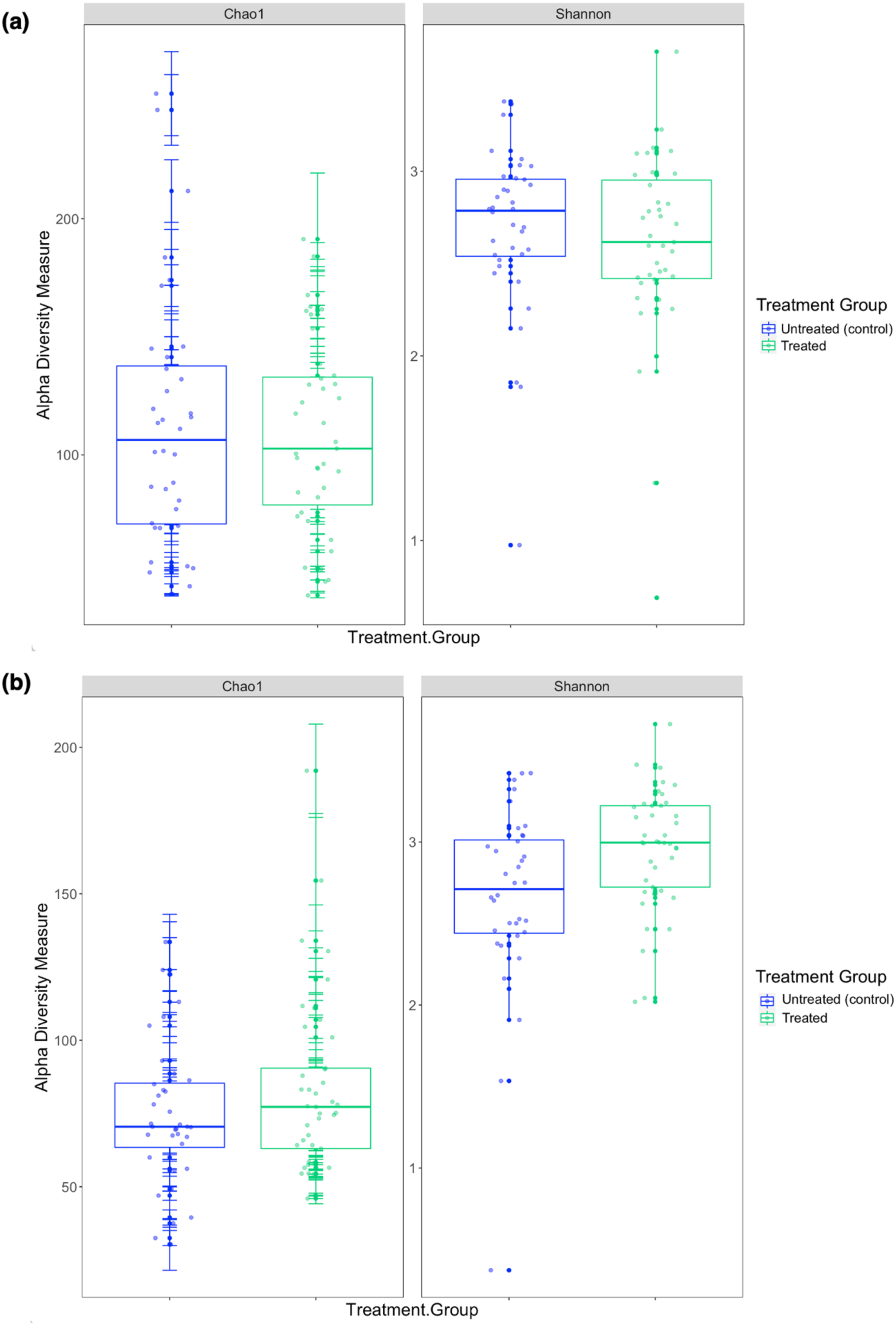
Comparison of the two alpha diversity measures, Chao1 index and the Shannon-Wiener index between treatment groups in the (a) 2019 breeding season and (b) 2020/21 breeding season.

The average diversity in the 2020/21 cohort was higher in the treated compared to the untreated (control) group (Table 2, Figure 5), which was reflected in the significant difference in the Shannon-Wiener index between treatment groups (p = 0.008). Richness was also, on average, higher in the treated group compared to the untreated (control) group (Table 2), however, there was no significant difference in the Chao1 index between treatment groups (p = 0.104). Similar results were observed in the 2020/21 breeding season, with no significant differences in alpha diversity between capture 1 and 2 in both the untreated (control) and treated groups for the Chao1 index (p =0.250 and p = 0.193) and the Shannon-Wiener index (p = 0.693 and p = 0.977).

When comparing alpha diversity between breeding seasons, there was a significant difference in both the Chao1 index (p = 0.001) and the Shannon-Wiener index (p = 0.017) between the 2019 and 2020/21 breeding seasons. On average, richness was greater in the 2019 cohort compared to the 2020/21 cohort while diversity was greater in 2020/21 (Table 2). Diversity and richness were also compared in treatment groups between breeding seasons. In the untreated (control) groups, there was no significant difference in diversity (p = 0.215) between seasons, however, there was a significant difference in richness (p = 0.013), with higher richness observed in the 2019 untreated (control) group (Table 2). There was a significant difference in both the Chao1 index (p = 0.025) and the Shannon-Wiener index (p = 0.001) between the treated groups between breeding seasons, with higher diversity in 2020/21.

### 3.2.2. Effect of treatment and pup morphometrics on alpha diversity

Two LMMs (models 1 and 3) were fitted to determine the effect of treatment and pup morphometrics on alpha diversity using the Chao1 and Shannon-Wiener indices. The first model tested differences between treatment groups and assessed the influence of treatment group, capture, pup age, weight (kg), standard length (cm) and pup sex.

When analysing Australian sea lion pups sampled during the 2019 breeding season as a cohort, there was a significant correlation between capture and Chao1 (R^2^=0.134, p=0.048) (Table S1). The richness of the gut microbiota fluctuated across captures and on average, was greatest in samples collected during capture 3 (100.87 ± 5.21). No significant correlations between any of the variables included in the first model and the Shannon-Wiener index were identified (Table S1). The correlation between pup morphometrics and alpha diversity was also investigated within each treatment group using the third LMM, which included the same factors as model 1 with treatment group excluded. In untreated (control) pups, there was no significant correlation between the Chao1 index and pup morphometrics, however, there was a significant correlation between capture and the Shannon-Wiener index (R^2^ = 0.176, p = 0.011) (Table S2). The mean diversity varied across captures and was highest after capture 3 (4.11 ± 0.38) before decreasing by capture 4 (3.74 ± 0.61). In Australian sea lion pups treated in the 2019 breeding season, there was no significant correlation between pup morphometrics and both the Chao1 index and the Shannon-Wiener index (Table S2).

In the 2020/21 cohort, there was a significant correlation between treatment group and both the Chao1 index (R^2^ = 0.107, p = 0.043) and the Shannon-Wiener index (R^2^ = 0.198, p = 0.002) (Table S1). On average, richness and diversity were higher in the treated compared to the untreated (control) group (Table 1). However, within the untreated (control) and treated groups there was no significant correlations between pup morphometrics and alpha diversity (Table S2).

### 3.2.3. Relationship between treatment, haematological parameters and alpha diversity

To evaluate the effect of topical treatment and haematological parameters on alpha diversity in the faecal microbiota of Australian sea lion pups, two LMMs (models 2 and 4) were fitted. The second model included treatment group, capture, pup age, hookworm status, TPP, TNCC and absolute leukocyte counts (neutrophil, lymphocyte, monocyte and eosinophil).

In the 2019 cohort, there was a significant correlation between capture and the Chao1 index (R^2^ = 0.134, p = 0.013) in the second LMM (Table S1); significant correlations between the Chao1 index and any of the other variables, including treatment group, were not determined. As with the results of model 1, there was no significant correlation with any factors and the Shannon-Wiener index. In the fourth model, which analysed the correlations between haematological parameters and alpha diversity within each treatment group, there were no significant correlations observed for the Chao1 index in either treatment group (Table S2). There was a significant correlation between capture and the Shannon-Wiener index in the untreated (control) group (R^2^ = 0.257, p = 0.019), while there were no significant correlations for the Shannon-Wiener index in the treated group. There was no significant correlation or relationship between haematological parameters and alpha diversity in either treatment group in the 2019 breeding season.

Analysis using the second fitted LMM found a significant correlation between treatment and both the Chao1 index (R^2^ = 0.135, p = 0.015) and the Shannon-Wiener index (R^2^ = 0.200, p = 0.04) in the 2020/21 breeding season (Table S1). Within the untreated (control) and treatment groups during the 2020/21 season there was no significant correlation between any factors included in model four and richness or diversity (Table S2).

## 3.3. Beta diversity

### 3.3.1. Beta diversity in the 2019 breeding season

Based on analysis of the differences in ASV abundance through the Bray-Curtis dissimilarity, 13.8% of the variation in the spread of the data could be explained by the first axis, and 8.7% could be explained by the second axis (Figure 6). There was no clear clustering of the data based on treatment group, indicating that dissimilarity between samples was not based on the treatment status of the individual (Figure 6). The results from the PERMANOVA models indicate that in pups sampled during the 2019 breeding season, pup ID was the most significant predictor of microbial similarity, and accounted for most of the variation in (Table 3). The capture event was also a significant predictor (p = 0.02), along with pup ID (p = 0.001) in the second model. The results from all models, presented in Table 3, determined that treatment group, hookworm status, age, pup weight, standard length and haematological parameters (TPP, TNCC, absolute neutrophil count, absolute lymphocyte count, absolute monocyte count and absolute eosinophil count; x10^9^/L), were all non-significant predictors of microbial similarity.

**Figure 6.**
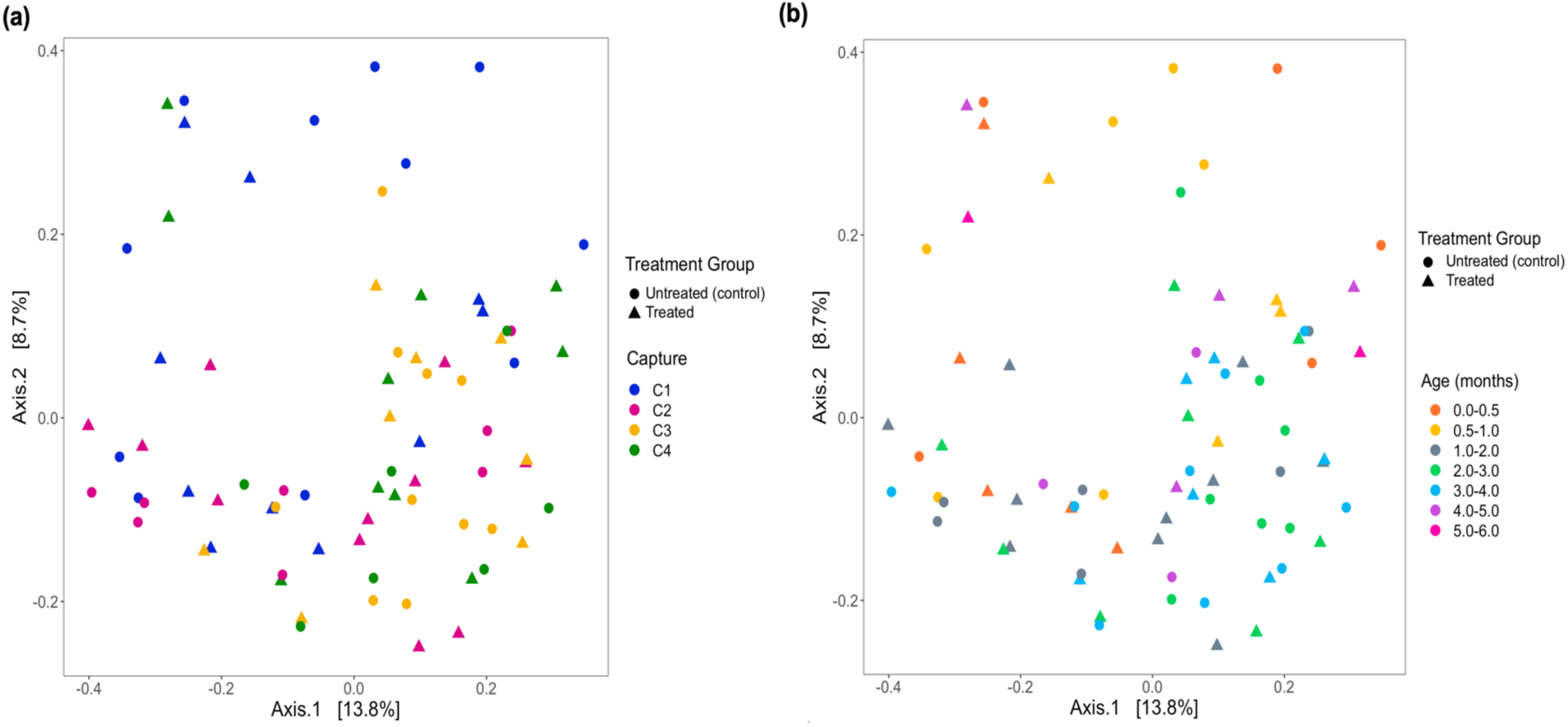
Beta diversity of faecal samples collected from Australian sea lion pups during the 2019 breeding season based on Bray-Curtis dissimilarities across (a) pup age groups (months) and (b) captures. In both plots, circles represent untreated (control) pups and triangles represent treated pups.

**Table 3.**
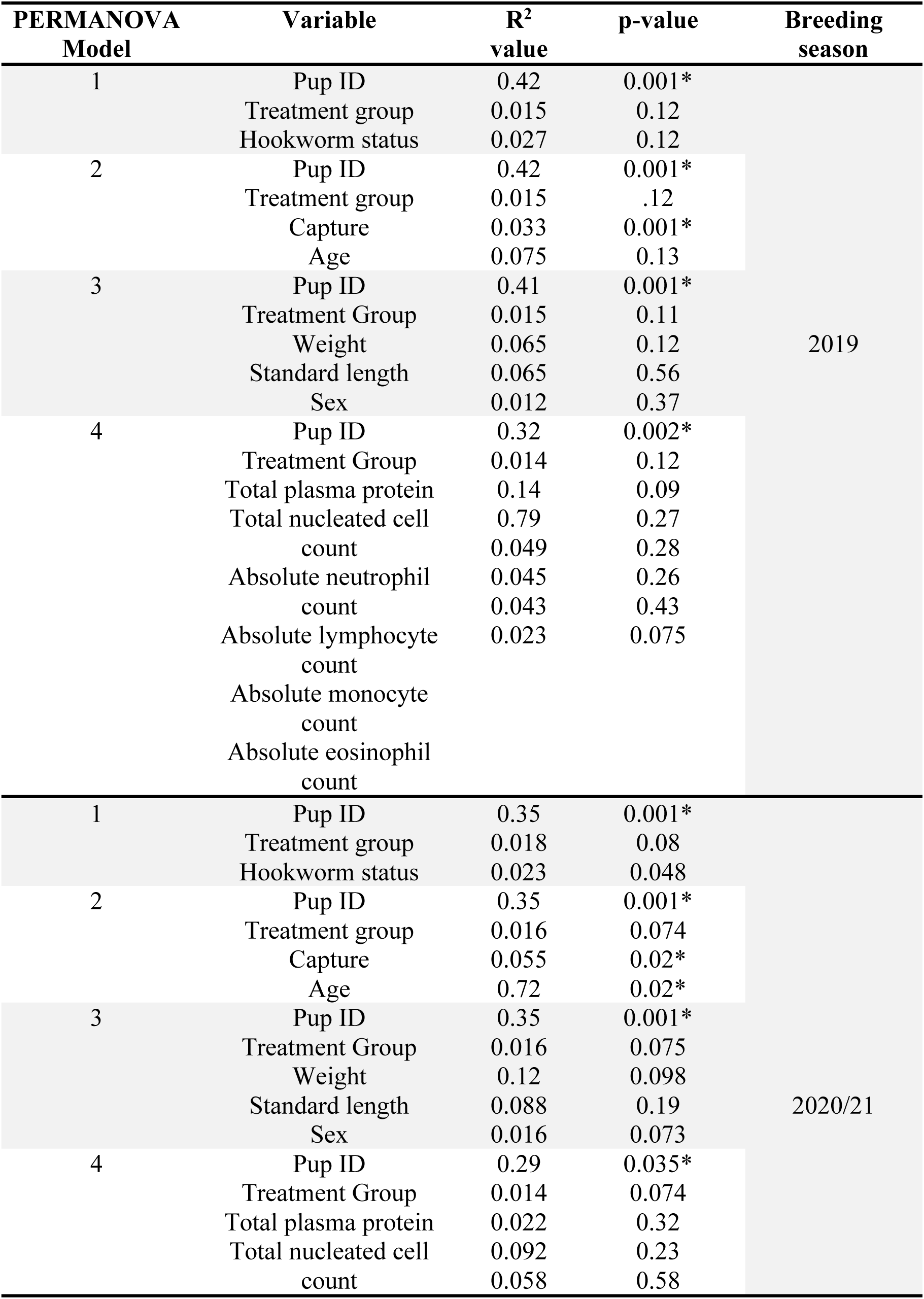
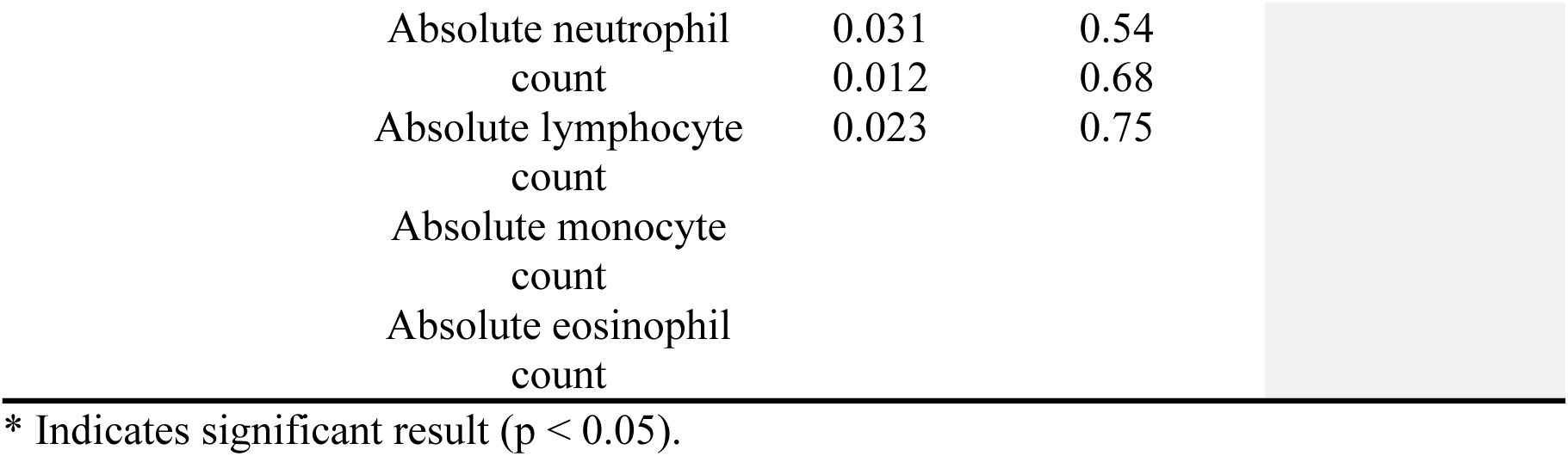
Results from PERMANOVA analysis from both 2019 and 2020/21 breeding seasons, with variables included in each model, variation accounted for by each variable (R^2^) and p-value.

There was a difference in the number of bacterial families identified between treatment groups, with five and seven in untreated (control) and treated pups, respectively. However, ANCOM analysis indicated that these unique features were not present at a high enough abundance to be detected as differentially abundant at the family or genus level between treatment groups.

### 3.3.2. Beta diversity in the 2020/21 breeding season

The analysis of the beta diversity of the faecal microbial community of pups sampled during the 2020/21 breeding season revealed that the first axis explained 13.1% of the variation while the second explained 7.6% (Figure 7). From the four PERMANOVA models, pup ID was the most significant predictor of similarity and accounted for most of the variation in each model (Table 3). In the second model, both pup age and capture were also significant predictors (p = 0.02). The contribution of capture and age to similarity can be visualised in the Bray-Curtis dissimilarity matrix, with samples collected at similar captures from pups of closer age groups clustering more closely (Figure 7).

**Figure 7.**
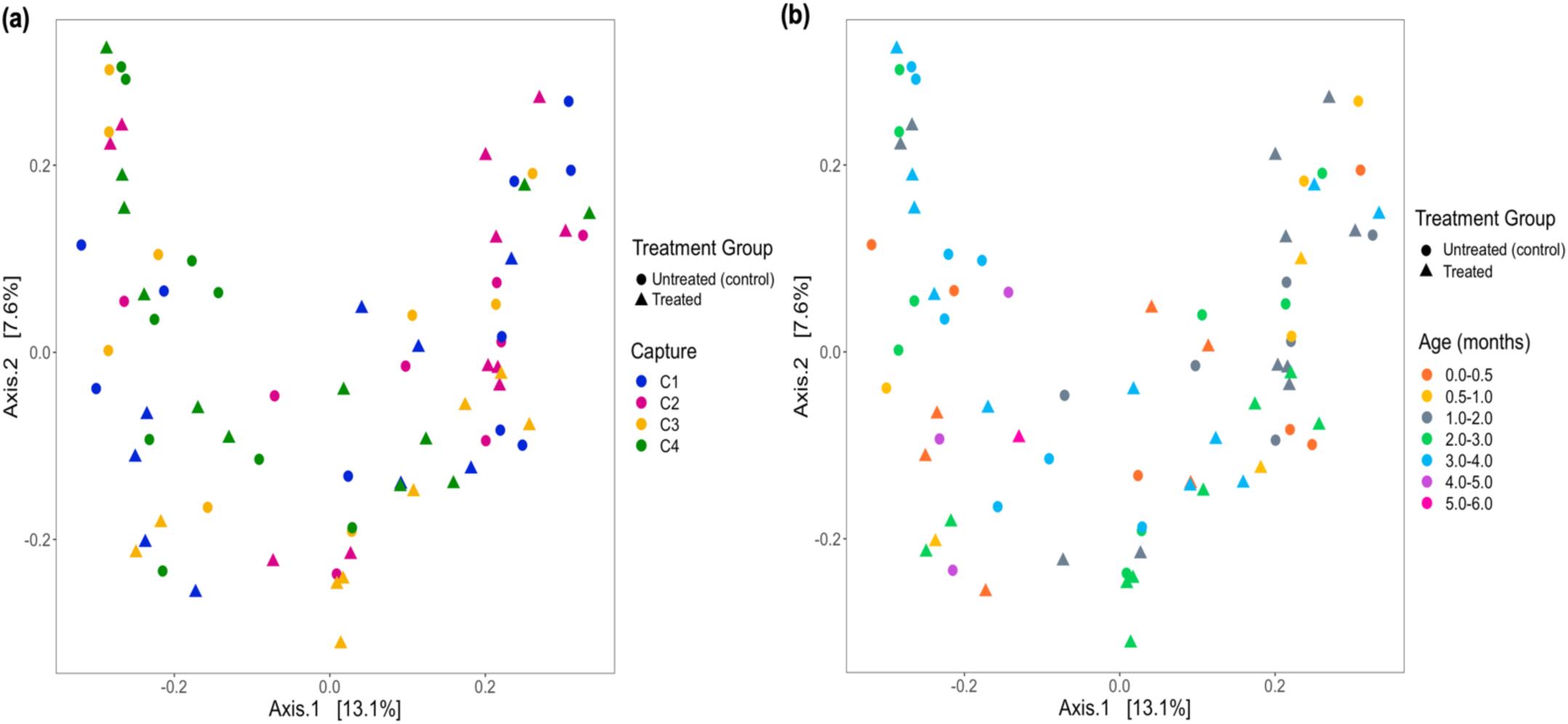
PCoA ordination of Bray-Curtis dissimilarities across (a) captures and (b) pup age groups (months), representing beta diversity of faecal samples collected from Australian sea lion pups during the 2020/21 breeding season. Circles represent untreated (control) pups and triangles represent treated pups.

In the 2020/21 breeding season, seven and five bacterial families were detected in samples collected from untreated (control) and treated pups, respectively. While there was a qualitative difference between the two groups, ANCOM analysis revealed that the features assigned to families only identified in untreated (control) pups were not present at a high enough abundance to be considered differentially abundant between treatment groups.

## 1.4. Discussion

This study aimed to determine whether topical ivermectin treatment and elimination of *U. sanguinis* infection altered the gut microbiota in endangered Australian sea lion pups. The findings from this study indicate that topical ivermectin treatment does not result in any compositional changes in the gut microbiota of Australian sea lion pups, suggesting that treatment does not cause any significant change to the functional capacity of the gut microbiome. This knowledge is crucial when considering the safety and efficacy of antiparasitic treatment in a free-ranging population, given the profound impact that the gut microbiota can have on host development and health through the regulation of the immune system, digestion and by protecting the host against pathogens (86–89).

It was hypothesised that treatment with topical ivermectin and resulting elimination of *U. sanguinis* would alter the gut microbiota of Australian sea lion pups when compared pre- and post-treatment. However, in the untreated (control) and treated groups in both breeding seasons, there were no significant differences in the composition of the gut microbiota between pre-treatment (capture 1) and post-treatment (capture 2) time points. For both treatment groups in both 2019 and 2020/21, the gut microbial communities were characterised by five bacterial phyla: *Fusobacteria, Firmicutes*, *Proteobacteria*, *Actinobacteria* and *Bacteroidetes*, and were dominated by *Fusobacteria*. These phyla have previously been identified as the main phyla in the gut of numerous pinniped species (28–35, 90). The relative abundance of each bacterial phylum differs between species and is likely influenced by differences in life histories, geographical ranges, environmental conditions, diet, and sampling techniques.

The outcome of helminth treatment on the gut microbiota of the host has been studied in humans, mice, and numerous wildlife species with varying results (51, 55–58, 91). Some human studies observed no effect between treated and untreated individuals pre- and post-treatment (55, 91, 92), while others found compositional changes and reduced alpha diversity in individuals’ post-treatment (56). In this study, the only significant difference in alpha diversity correlated with treatment group was observed in Australian sea lion pups treated during the 2020/21 breeding season. Both the Chao1 and Shannon-Wiener indices were higher in treated pups compared to untreated (control) pups. However, in treated pups, there was no significant difference in alpha diversity between pre- and post-treatment time points (capture 1 vs. capture 2), suggesting that the removal of hookworm infection after treatment is not the cause of the significant difference observed. Furthermore, hookworm status was not associated with significant changes in alpha diversity in either of the treatment groups.

Despite differences in richness and diversity of microbial community in Australian sea lion pups, there were no significant differences in microbial community composition between treatment groups, which is attributed to the topical method of administration. In Amur tigers (*Panthera tigris altaica*) treated with oral Fenbendazole and ivermectin, significant changes in both the gut microbiota and faecal metabolic phenotypes were seen, suggesting that treatment both disturbed the microbial community within the gut as well as metabolic homeostasis (58). Significant differences in gut microbial composition and diversity were also observed in Asian elephants (*Elephas maximus*) administered oral Albendazole treatment (57) and in humans treated for helminth infection with oral ivermectin (56).

Parasite presence in the gastrointestinal tracts of humans, livestock and wildlife species have been found to have differing impacts on gut microbial diversity, largely dependent on host and parasite species (47, 52, 53). In the present study, there was no clear association between the presence of *U. sanguinis* and taxonomic assignment of ASVs or microbial diversity in both treatment groups regardless of breeding season. As *U. sanguinis* infection is endemic in the Australian sea lion population (13, 14), it is possible that the gut microbiota of Australian sea lions has co-evolved with this parasite and is not influenced by the presence or absence of *U. sanguinis*, either through natural clearance due to age-related parasite senescence or host immune response, or due to parasitic treatment.

In pinnipeds, the relative abundance of *Fusobacteria* appears to be highest in pups and decreases with age (28, 30, 31). For example, in southern elephant seals, there was a significantly higher abundance of *Fusobacteria* in pups compared to sub-adults and adults (31), and the relative abundance of *Fusobacteria* has been found to decrease steadily during the weaning period of Pacific harbor seals (28). Changes in bacterial phyla abundance was also reported with age in Australian fur seals, with significant shifts observed between pup, juvenile and adult life stages (32). In Australian sea lion pups, a shift in the relative abundance of bacterial phyla and families across capture events was observed, although this shift is likely an age-related change, with up to five months between first and fourth captures. The results from the Bray-Curtis dissimilarity matrix suggested that the gut microbiota was more similar in pups of the same age, regardless of treatment group. A higher relative abundance of *Fusobacteria* was also observed in the current study in both treatment groups between the 2019 and 2020/21 breeding seasons, where *Fusobacteria* was the most abundant bacterial phyla. Bacterial families belonging to the *Fusobacteria* phylum were also the most dominant in the gut microbiota of Australian sea lion pups between both breeding seasons.

The trends observed in the 2020/21 breeding season also suggest that the relative abundance of *Fusobacteria* decreases with age, with the highest relative abundance occurring in samples collected during the first capture and steadily decreased across subsequent captures, irrespective of treatment group.

The unique 18-month breeding cycle of the Australian sea lion (93, 94) means that the 2019 breeding season at Seal Bay occurred during the Austral winter, while 2020/21 began during summer. As such, it was expected that seasonal and environmental variations could influence the gut microbiota of Australian sea lion pups. There were differences in the relative abundance of the more abundant phyla, *Fusobacteria*, *Firmicutes*, and *Actinobacteria*, and in two bacterial families that were present in both seasons, *Fusobacteriaceae* and *Clostridiaceae*. Within the 2019 and 2020/21 breeding seasons, there were no significant changes in the relative abundance of bacterial phyla or families across captures. However, qualitative changes in microbial community composition were observed; in 2019, the relative abundance of *Fusobacteria* remained relatively stable across all captures in both treatment groups, while the abundance of *Firmicutes* steadily decreased, and *Actinobacteria* and *Proteobacteria* increased. The relative abundance of *Fusobacteria* decreased more markedly in pups sampled during 2020/21 and was accompanied by a decrease in *Firmicutes,* and an increase in *Proteobacteria*. This was also reflected at the family level, with the relative abundance of *Fusobacteriaceae*, a family belonging to the *Fusobacteria* phylum, decreasing from capture 1 to capture 4 in both treatment groups. Despite the significant differences between breeding seasons, the gut microbiota of Australian sea lion pups was dominated by the same bacteria phyla in both treatment groups in 2019 and 2020/21, suggesting that the functional capacity of the gut microbial community is similar across treatment groups and breeding seasons. When comparing richness and diversity of the gut microbiota between seasons, there was again a significant difference; the richness of the gut microbiota was higher in Australian sea lion pups sampled in 2019, while diversity was, on average, higher in the 2020/21 cohort. The richness and diversity of the gut microbiota was significantly different in the treated group when compared between 2019 and 2020/21, and only richness differed significantly in the untreated (control) groups between seasons. The significant differences observed between seasons highlights the influence of external environmental conditions on the richness, diversity, and composition of the microbiota in Australian sea lion pups. Further investigations are required to determine the specific environmental characteristics contributing to these differences.

Host factors including pup weight, standard length, sex, and haematological parameters were also investigated to elucidate their relationship with treatment and the gut microbiota. In pups from both treatment groups sampled in the 2019 and 2020/21 breeding seasons, weight, standard length, and sex of the pup did not significantly influence the gut microbial composition. In sexually dimorphic pinniped species, differences in relative abundance of bacterial phyla and the relative abundance of ASVs or OTUs have been observed between male and females (28, 31). In adult southern elephant seals, differences between sexes were attributed to differences in diet rather than body mass (31). When investigating the contribution of sex to the gut microbiome in weaned southern elephant seal pups (one to three months old), Stoffel et al. (2020) (28) found differences between sexes even though the male and female pups were indistinguishable, suggesting that some gut microbes are sex-specific and necessary for future adult feeding strategies. While Australian sea lions are a sexually dimorphic species, Australian sea lion pups sampled as part of this study were a maximum of four to six months old and completely dependent upon their mothers for nutrition. Furthermore, Australian sea lion pups are not weaned until 18 months of age (2). As a result, sampled pups were likely too young for any sex-associated differences in microbial composition to occur. The haematological parameters of Australian sea lion pups can be used as indicators of overall health and improvement of total leukocyte counts have previously been associated with ivermectin treatment in previous studies (13, 18, 19). However, there was no significant association between TPP, TNCC, total leukocyte, neutrophil, lymphocyte, monocyte and eosinophil counts and alpha diversity, beta diversity, or the relative abundance of any bacterial phyla or family.

## 1.5. Conclusion

Topical ivermectin treatment of Australian sea lion pups has been identified as a potential management strategy to improve the health of free-ranging pups in a declining population (19), however, unexpected consequences of treatment need to be explored to ensure deleterious outcomes do not ensue. Characterisation of the gut microbiota of free-ranging untreated (control) and treated Australian sea lion pups revealed that topical ivermectin treatment and elimination of *U. sanguinis* infection did not alter the composition of the gut microbial community. Despite some minor differences, the absence of statistically significant alterations to the composition of the gut microbial community, together with the known benefits of ivermectin treatment for Australian sea lion pup health and growth, suggest that topical ivermectin could be an effective management strategy to mitigate the impact of endemic hookworm disease on population growth in this species.

## Acknowledgements

We thank the staff at Seal Bay Conservation Park, Kangaroo Island, Department of Environment and Water (DEW), South Australia, particularly Melanie Stonnill, Melanie Stephens, Alana Binns and Tanya Rosewarne for field assistance and logistical support; Simon Goldsworthy and the South Australian Research and Development Institute (SARDI) for field assistance. We would also like to thank Scott Lindsay, Shannon Taylor and Matthew Gray for assistance in sample collection and Janet Simpson for logistical support.

---

**Supplementary Table 1.**
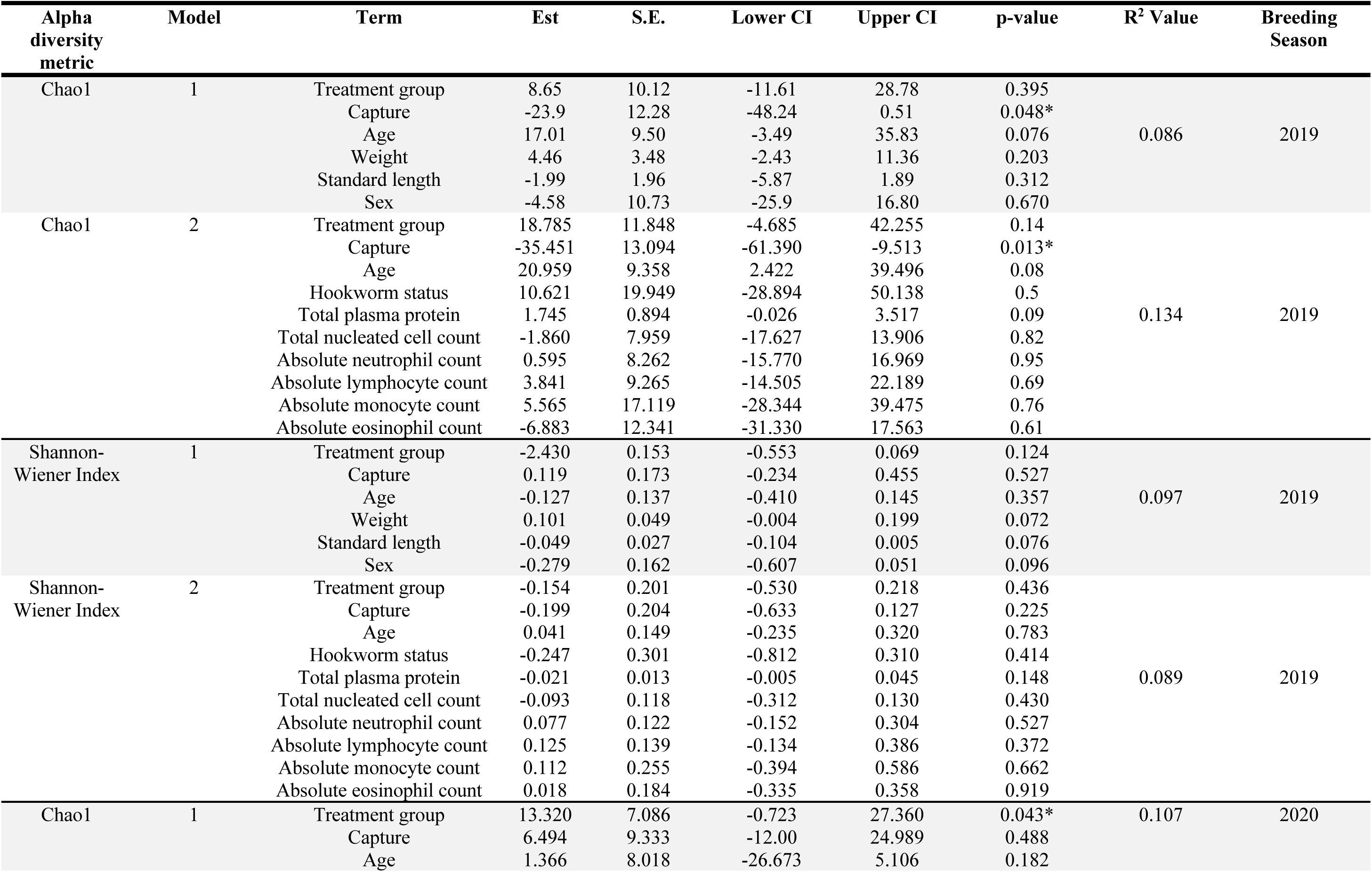
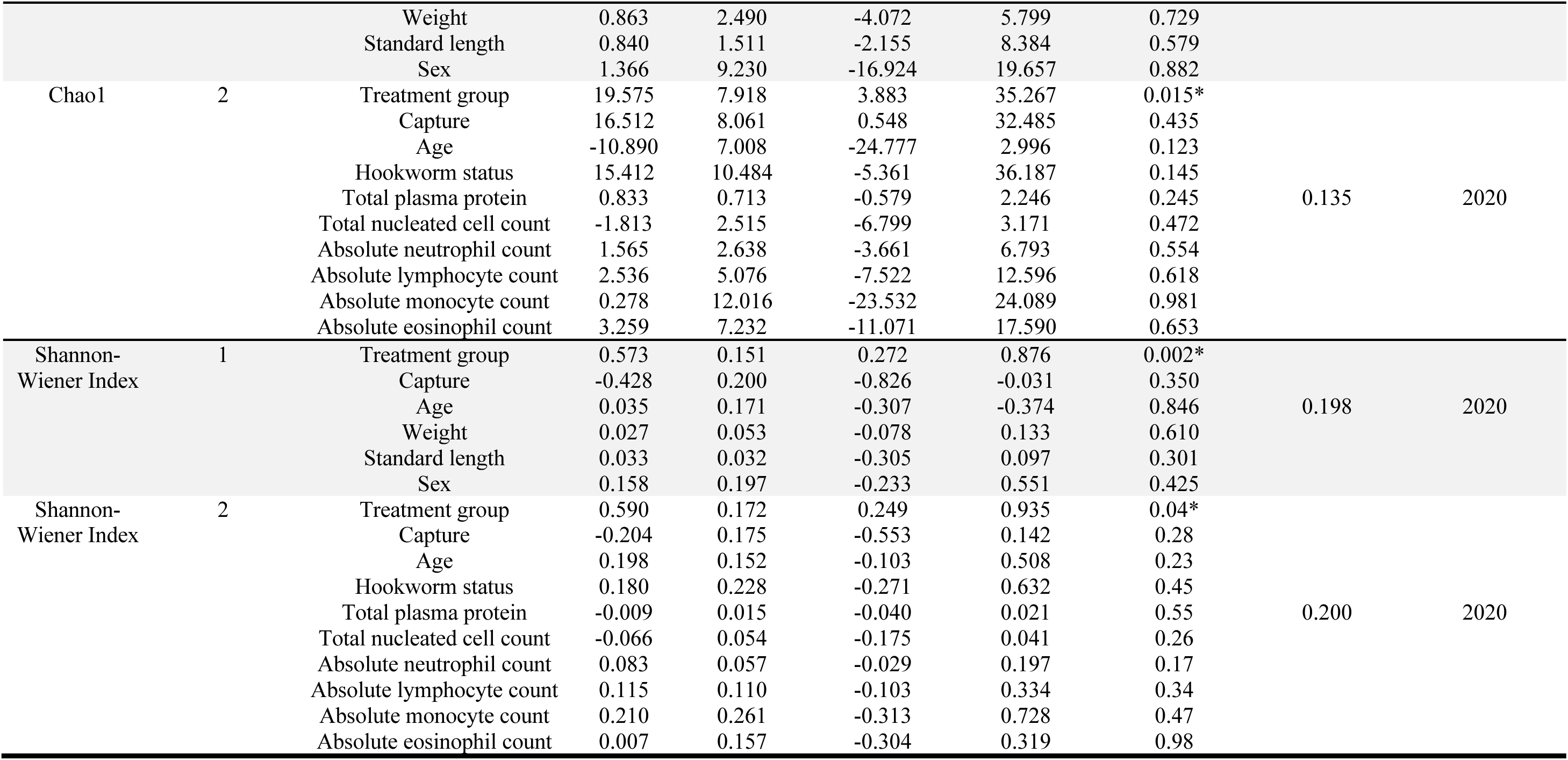
Results from linear mixed models one and two, analysing the correlations between pup morphometric and haematological data and alpha diversity in the 2019 and 2020/21 breeding seasons.

**Supplementary Table 2.**
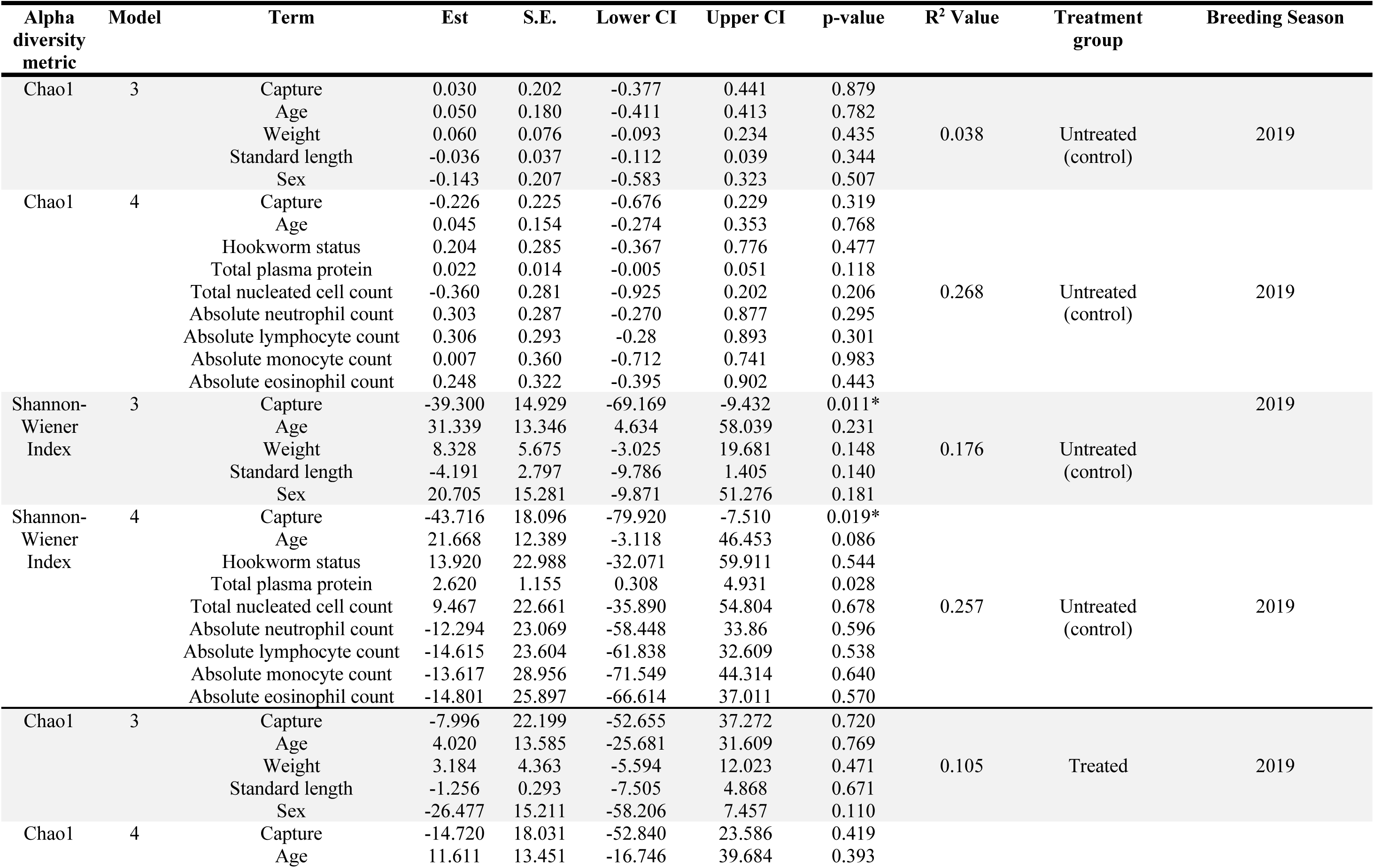
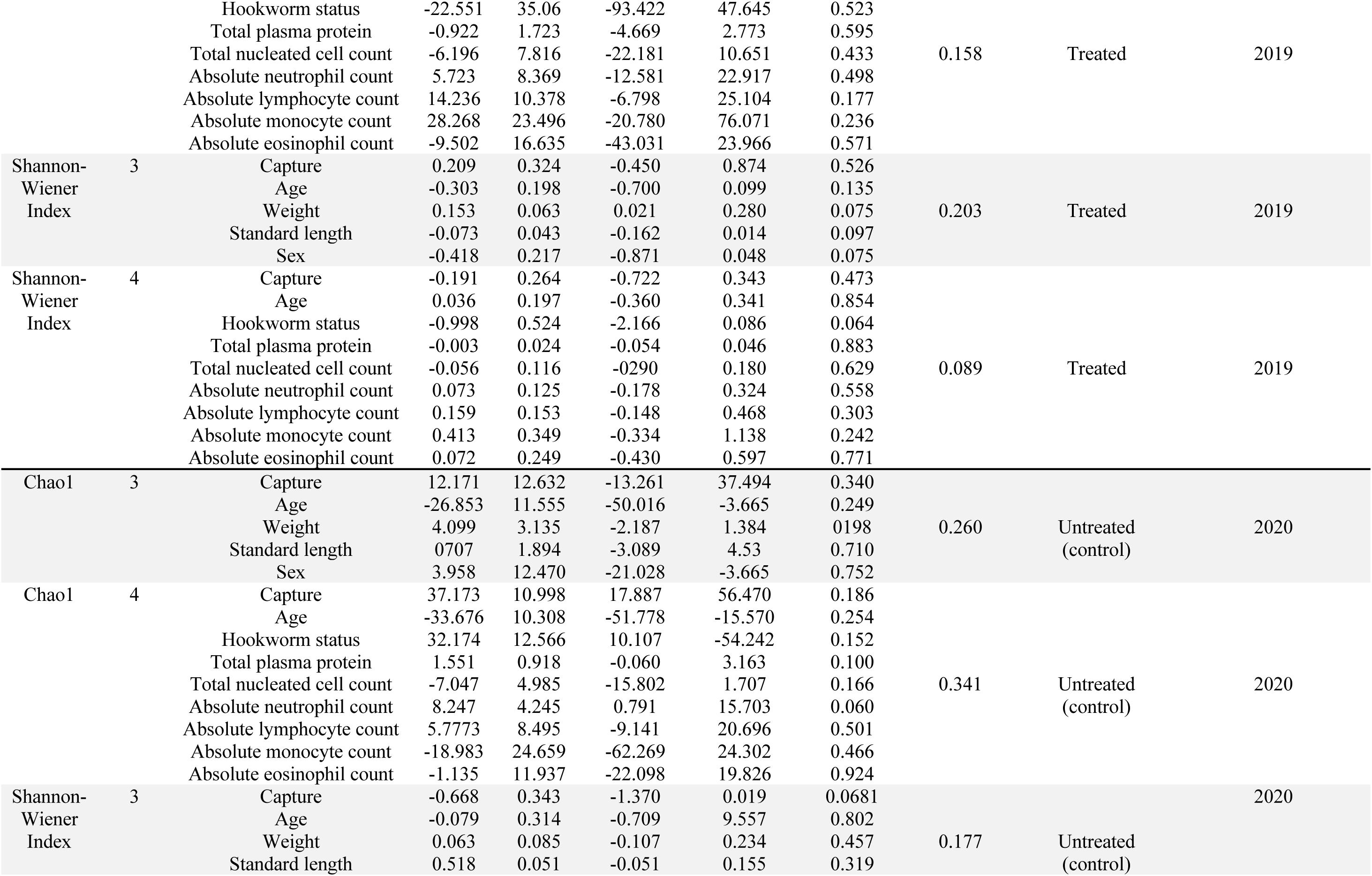
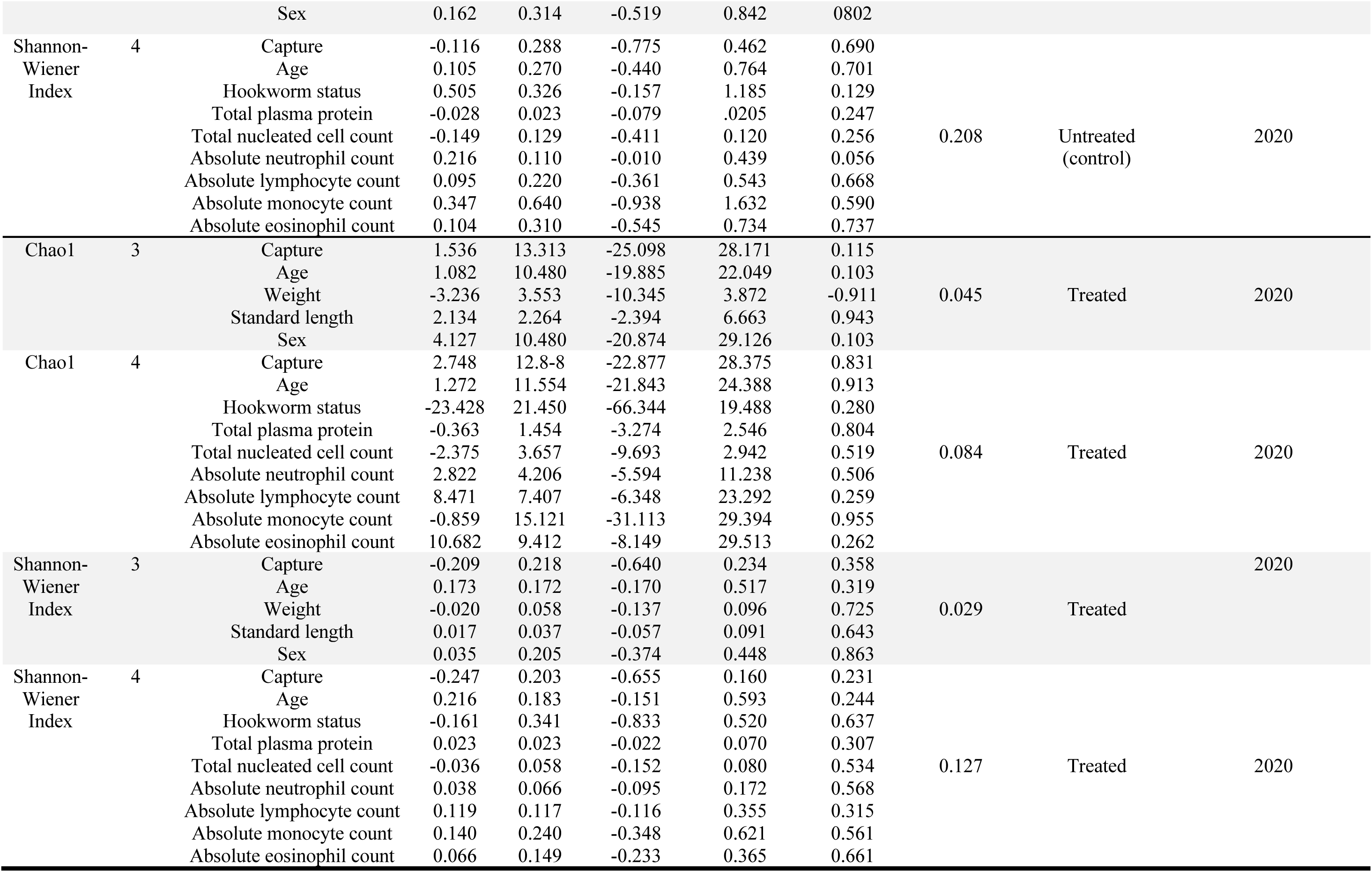
Results from linear mixed models three and four, analysing the correlations between pup morphometric and haematological data and alpha diversity within each treatment group in 2019 and 2020/21.

